# Branching and extinction in evolutionary public goods games

**DOI:** 10.1101/2020.08.30.274399

**Authors:** Brian Johnson, Philipp M. Altrock, Gregory J. Kimmel

**Affiliations:** Department of Integrated Mathematical Oncology, H. Lee Moffit Cancer Center and Research Institute, Tampa, FL 33612, USA

## Abstract

Public goods games (PGGs) describe situations in which individuals contribute to a good at a private cost, but others can free-ride by receiving their share of the public benefit at no cost. PGGs can be nonlinear, as often observed in nature, whereby either benefit, cost, or both are nonlinear functions of the available public good (PG): at low levels of PG there can be synergy whereas at high levels, the added benefit of additional PG diminishes. PGGs can be local such that the benefits and costs are relevant only in a local neighborhood or subset of the larger population in which producers (cooperators) and free-riders (defectors) co-evolve. Cooperation and defection can be seen as two extremes of a continuous spectrum of traits. The level of public good production, and similarly, the neighborhood size can vary across individuals. To better understand how distinct strategies in the nonlinear public goods game emerge and persist, we study the adaptive dynamics of production rate and neighborhood size. We explain how an initially monomorphic population, in which individuals have the same trait values, could evolve into a dimorphic population by evolutionary branching, in which we see distinct cooperators and defectors emerge, respectively characterized by high production and low neighborhood sizes, and low production and high neighborhood sizes. We find that population size plays a crucial role in determining the final state of the population, as smaller populations may not branch, or may observe extinction of a subpopulation after branching. Our work elucidates the evolutionary origins of cooperation and defection in nonlinear local public goods games, and highlights the importance of small population size effects on the process and outcome of evolutionary branching.

## 1. Introduction

Emergence, evolution and persistence of subpopulations with differing traits or strategies remains an open question for many systems in ecology and evolution [1–3], cancer biology [4], and human social interactions [5, 6]. Speciation events in evolution are difficult to observe because they occur over long timescales [7, 8]. In contrast, somatic evolution occurs over shorter timescales, but has also proven to be difficult to understand due to complications arising from limited sampling ability [9–12]. Evolutionary game theory can be used to explore and explain the emergence and disappearance of traits or strategies as the environment and the population composition change over time [13]. Branching and speciation in evolutionary population games can be studied using adaptive dynamics, which allows us to study evolutionary branching [14], coexistence, and extinction events in continuous trait or strategy space over time [15, 16]. For example, this framework has been used to study branching of complex behavioral traits [17], polymorphism in cross-feeding [18], and the emergence of cooperators and defectors in social dilemma games [19].

A well-known example of a social dilemma among groups of individuals is the “Tragedy of the Commons” [20, 21], which emerges when individuals in a group contribute to a common good that is shared independent of contribution. All individuals tap into a benefit or common resource, e.g. via consumption. As the resource is depleted by the cumulative consumption of all individuals, the benefit to each individual erodes, until there is very little to none left. Individuals would have been better off in a state with moderate consumption, against the rational choice to maximize their own consumption. In this example, one might call those who maximize their consumption of the resource “defectors”, due to the detrimental effect their actions have on others. Such dilemma situations are often found to describe aspects of interactions among cancer cells [4, 22], banded mongooses [1], prairie dogs [23], predator inspection in fish [24], or mobbing of hawks by crows [25]. Microbes provide public goods by secreting useful chemicals [26, 27]. Yeast cells synthesize and spill essential nutrients into their surroundings [2], and were observed to coevolve [28]. Such potentially complex interactions can strongly influence the eco-evolutionary dynamics between cooperators (public good producers) and defectors (free-riders). However, it has less often been examined how these two distinct strategies emerge in the first place, which can be answered by studying the adaptive dynamics, or evolutionary invasion analysis [15, 29].

A key assumption in models of public good (PG) dynamics is the effect of the PG on the payoff or fitness of an individual [30]. There can be diminishing returns that naturally leads to non-linear benefit functions [31]. Examples include linear, convex, concave, and sigmoidal benefit as a function of the available PG [4, 30, 32–38]. Here, we consider a sigmoid relationship between the level of PG and the resulting benefit, as well as a nonlinear cost of production. We are interested in the evolution of two key traits that change these relationships. First, we consider the size of the group among which the public goods game is played, also called the neighborhood. Second, we consider the level of PG production of each individual. We are interested in the evolution of the population in this two dimensional trait space. Intuitively, it might be clear that cooperators who exhibit high levels of production favor small neighborhoods, whereas defectors with low levels of production can only survive if their neighborhood sizes are sufficiently large. It has been unclear so far how these distinct strategies can emerge from initially homogeneous populations, and what role small population sizes play in the extinction of an entire subpopulation.

We employ adaptive dynamics [15] to show how an initially monomorphic population in trait space can evolve into two distinct subpopulations comprised of cooperators and defectors. When the population size is large, the trait evolution is deterministic and easier to predict. However, when the population size is small, stochastic fluctuations become relevant. We seek to understand how small populations behave and how the results deviate from those predicted by the deterministic theory. In recent years, investigation of the impact of small populations has been studied, most notably in the work of Wakano and Iwasa [39], Claessen et al. [40], and Debarre and Otto [41]. These previous works have revealed that branching in stochastic systems is either delayed or does not occur, and extinction of certain types may occur following branching. While most of the previous work focuses on games with a single evolving trait, our work explores the impact of branching and extinction in a two-dimensional game. Because many relevant processes begin with small populations, further exploration of co-evolving traits in finite populations is warranted.

This manuscript is structured as follows. In Section 2, we introduce the evolutionary population game, how it proceeds over time, and the methods we use to analyze the results. In Section 3, we present our simulation results and compare them to analytical approaches that predict trait evolution. In Section 4, we summarize our findings, put them into context, and highlight interesting areas for future work.

## 2 Methods

### 2.1 Model Introduction

Benefits that are produced and shared by a focal individual are often only available within a finite subset, or neighborhood [42, 43]. Furthermore, some individuals may interact more locally than others. Here, we are interested in how production and neighborhood size evolve when they become part of the selection process.

Models of evolutionary public goods games often assume a linear relationship between benefits and the number of cooperators [44, 45]. Yet the collective benefit of a public good may be nonlinear [32, 37, 46, 47]. There may be increasing returns when public good increases from low to higher levels, and diminishing returns when it levels further increase from high to very high levels (i.e. saturation), leading to a sigmoidal public good to payoff relationship.

The nonlinear payoff function of our model was previously introduced in Kimmel et al. [37], who considered a sigmoidal benefit proportional to the amount of public good shared among a fixed neighborhood of individuals, and a fixed cost for production. Neighborhood size, *n*, determines the subset of individuals among which the focal individual shares the PG. The population size, *N*, is fixed throughout the process, and we consider evolution in steps of generations between which the entire population is replaced by their offspring population, in which relative abundances change due to selection and randomness.

To study adaptive dynamics, we model public good production as a continuous trait, and use previously established nonlinear benefit function [37]. We also introduce a sigmoidal cost function, and allow neighborhood size to evolve as a second trait. The population game works as follows. Consider a focal individual with production level *y* and a neighborhood of size *n_y_* (of which the individual itself is a part). Then, by any probabilistic assembly, there are *n −* 1 neighbors, their respective levels of production can be labelled *x_i_* (*i* = 1,…, *n_y_* − 1). The public good production-traits, *y* and *x_i_*, are continuous variables between 0 and 1. In the evolutionary process the neighborhood size-trait is also a continuous variable between 1 and *N*.

When the payoff is calculated, the neighborhood size of the focal individual, *n_y_*, is rounded to a discrete integer between 1 (private good) and the population size, *N*. For each focal individual in each generation, *n_y_ −* 1 neighbors are drawn from hypergeometric sampling of the population, excluding the focal individual. For further details, see Appendix A. We can then calculate the focal individual’s available public good, *G* as the result of its own and its neighbor’s production

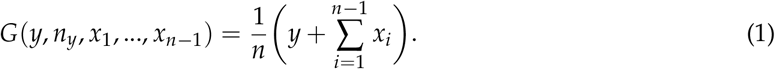

If **x** represents the chosen neighbors, *x_i_* (*i* = 1,…, *n_y_ −* 1), then the nonlinear (sigmoidal) benefit to the focal individual is [37]

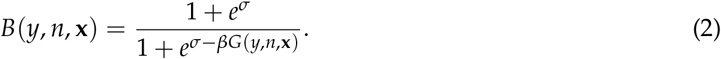

The benefit function can be altered by adjusting *β*, which controls for the PG-dependent benefit, and *σ*, which controls for the PG-independent benefit. Unless otherwise specified, we set *σ* = 2 and *β* = 5 (see Table 1), as we have shown that these parameters should, in principle, allow for coexistence [37].

**Table 1:** The table shows the parameters and the typical values we used running the game. Some were taken from previous literature (as referenced) with similar games, while others were chosen to fit the game dynamics.

| Parameter | Meaning | Typical Value | Reference |
| --- | --- | --- | --- |
| $\sigma$ | Scales PG-independent benefit | 2.0 | [37] |
| $\beta$ | Scales PG-dependent benefit | 5.0 | [37] |
| $\kappa$ | Cost magnitude | 0.5 | [37] |
| $\omega$ | Cost shape | 2.0 | this work |
| $\mu_x$ | Probability of mutation in individual production | 0.01 | [19] |
| $\mu_n$ | Probability of mutation in individual NS | 0.01 | this work |
| $s_x$ | Production mutation standard deviation | 0.005 | [19] |
| $s_n$ | Neighborhood size mutation standard deviation | 0.2 | this work |

Cost of public good production, *C*(*y*) is determined only by the production of the individual. It is controlled by *κ*, which specifies its relative magnitude, and *ω*, which specifies the shape

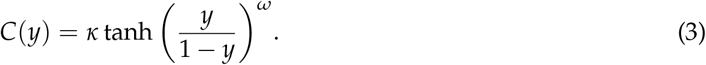

Typically, the cost function parameters were chosen to replicate the general sigmoidal shape of the benefit function, saturating at high production, but also increasing slowly for low production, with the largest slope for medium production.

Using benefit and cost functions, we can determine the payoff to the focal individual characterized by production level *y*:

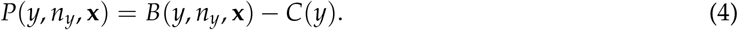

Overall, this form makes it clear that the optimal strategy of one trait depends on individual context.

The population is initially seeded in production and neighborhood trait space via two independent and normally distributed distributions. This monomorphic population evolves over time in trait space. In certain scenarios the population can branch and lead to two distinct subpopulations (Figure 1 A).

**Figure 1:**
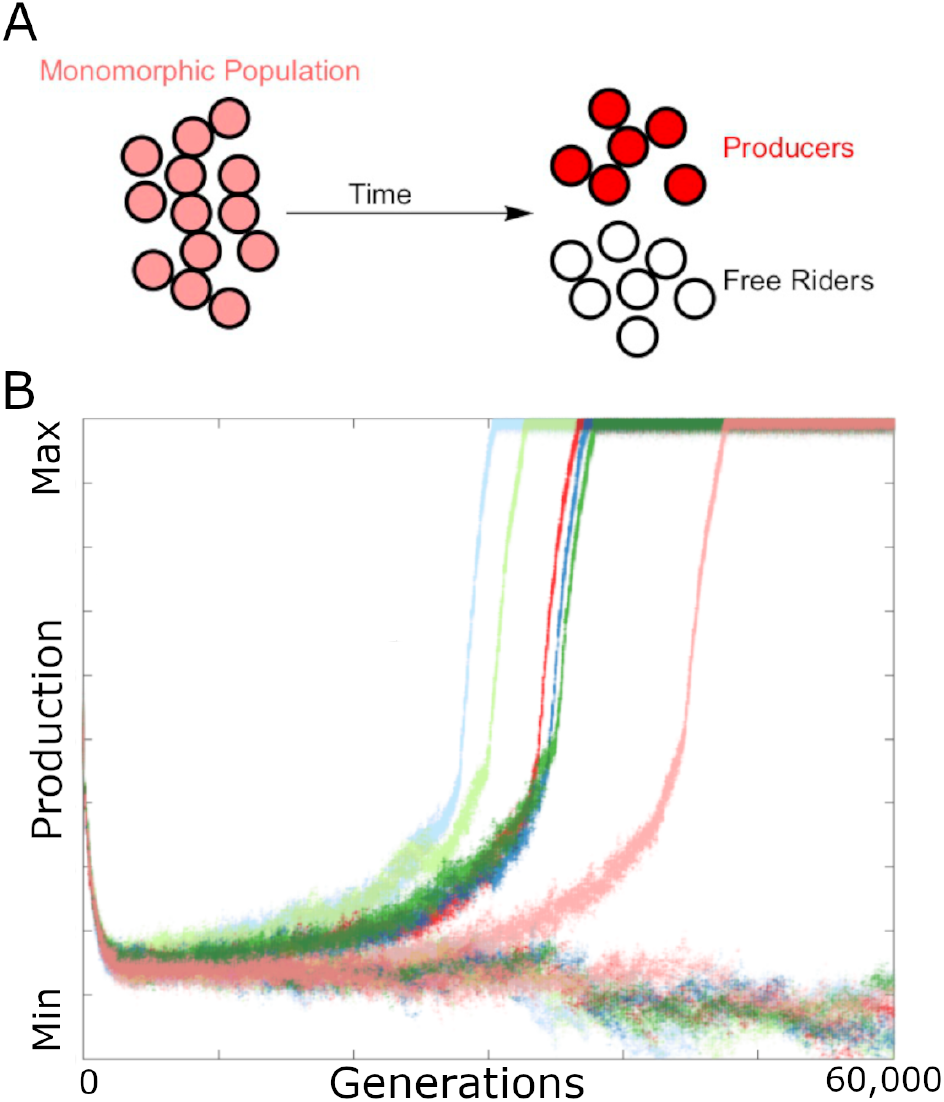
Model Introduction. **A:** A schematic view of an initially monomorphic population branching into two distinct subpopulations, with color indicating trait. **B:** An example of the stochastic branching process, showing production over time for a population size of 500. Six simulations with identical initial conditions are shown, with color differentiating simulations. All of the simulations branch, as predicted by deterministic theory, though the generation at which branching occurs varies. These simulations all use default parameters: *β* = 5, *σ* = 2, *κ* = 0.5, *ω* = 2 (table 1).

In our model, trait evolution follows the process proposed by Doebeli et al. [19]. Time advances in discrete generational time steps during which each individual’s payoff is calculated. After a focal individual interacts with neighbors to determine its payoff, the payoff is then compared to the payoff of second randomly selected individual, which we call the opponent. The opponent has some probability to replace the focal individual. Replacement probability is normalized by *α*, which is equal to the largest payoff gap in the current generation

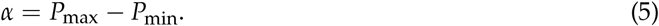

Then, with probability *q*, the opponent replaces the individual and becomes the parent of an offspring in the next generation:

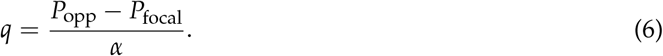

If replacement does not occur, the focal individual is the parent. The offspring will have the traits of the parent, possibly altered by mutation.

The mutation process involves the two probabilities of mutation, *μ_x_* and *μ_n_*, (see table 1), such that on average, one out of every 1/*μ_x_* individuals is expected to mutate in a given generation for production and one of every 1/*μ_n_* is expected to mutate in neighborhood size. Mutations occur independently in each trait, and are distributed normally with the parent’s trait as the mean and a chosen standard deviation (*s_x_* or *s_n_*). For a full treatment of the game details and pseudo-code, see Appendix A.

### 2.2 Estimation of branch location and probability of branching

We are interested in branching events in trait space. To determine the branch location in time and in trait space, we utilized a *k*-means supervised clustering algorithm (as implemented in MATLAB R2019b, with *k* = 2) in the following way. At a branch event, the population splits into two groups. A branch can thus be defined to occur when the means were separated by a chosen threshold. For the purposes of evaluating branching, we set a threshold of 0.2 in the production trait. Using this approach, we obtained numerical results for the probability of branching and the generation to branch (if it occurs) for various population sizes (see Appendix B for details).

Our two-dimensional trait space differs from the one-dimensional system in an important way; we have a finite window in trait space at which branching can occur. The population, while still monomorphic, drifts along the “branching curve”, a line of attractor points (solid black line in Figure 2 A, C, E), along which branching is favored. Once the population sufficiently decreases in neighborhood size-trait, the population moves quickly to maximum production and minimum neighborhood size, such that branching is no longer possible. Hence, there is a finite time interval during which the population remains in a region favored to branch. In simulations, branching or no branching occurs within a finite amount of time.

**Figure 2:**
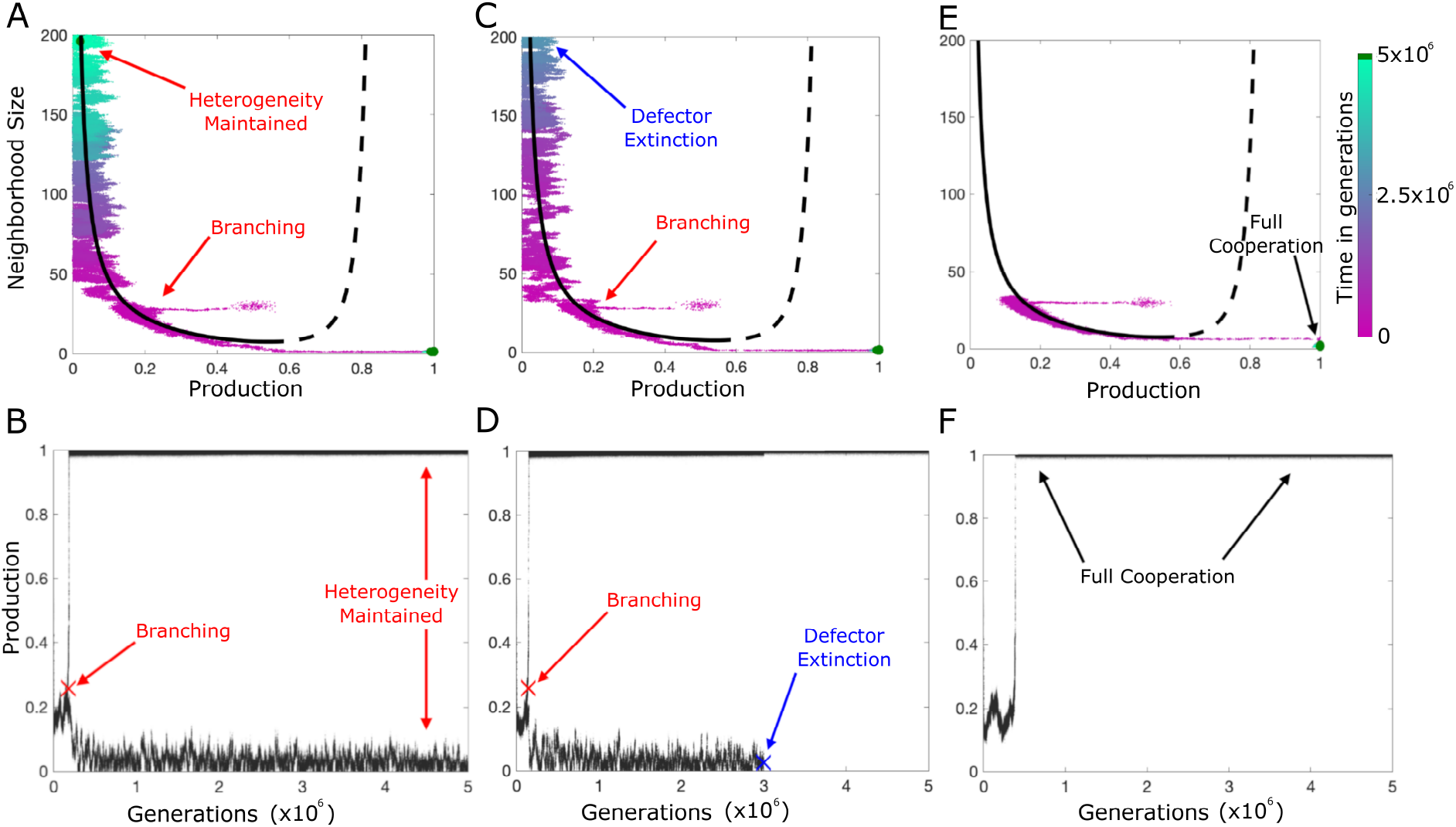
Long-term outcomes with the same population size. The upper panels show a trait space view, with the solid and dashed black line indicating the equilibrium points at which the selection gradient is zero for a monomorphic population. Solid represents the attracting set, and dashed represents the repulsive set. The lower panels show the corresponding temporal view in the production trait. In the upper panels, color indicates generation, with dark green indicating the traits of the final generation. Initial states of all populations are normally distributed with production of 0.5 and a neighborhood size of 30. “Default” parameters (*β* = 5, *σ* = 2, *κ* = 0.5, *ω* = 2) were used for all simulations, and all simulations contain 200 individuals run for 5 million generations. **A & B:** Branching occurs, and is maintained, as noted by the two separate groups of dark green at the final generation. Red ‘X’ in **B** denotes branching point. The final result is a heterogeneous population. **C**& **D:** Branching occurs, but extinction of the free-rider subpopulation occurs before the final generation, as noted by the lack of a dark green group in the upper left. Again, red ‘X’ in **D** denotes branching point. Additionally, blue ‘X’ denotes extinction point. The final result is a monomorphic population of producers, which cannot branch again. **E**& **F:** Branching never occurs, and will never occur. The population moves out of the region where branching is possible. The result is also an all-producer population.

In contrast, single trait adaptive dynamics, such as those studied by Doebeli et al. [19], Wakano and Iwasa [39], and Debarre et al. [41], have populations which are favored to branch for as long as the simulation runs. In these systems it becomes difficult to calculate the probability that branching occurs. Our two-dimensional system does not have this problem, as we can say with certainty whether branching occurred or not within a finite window. The impact of this window will be discussed further, as it could have interesting consequences on the overall evolutionary dynamics.

## 3. Results

In this section, we assess the successes and failures of adaptive dynamics as it applies to our game. First, we use the deterministic approach to make predictions under the assumption of an infinitely large population. Following these predictions, which we know are inaccurate for small population sizes, we discuss why they fail and how we can adapt our predictions to capture the stochastic fluctuations that become important in small populations.

### 3.1 Monomorphic populations tend toward the branching curve

The payoff function and the defined rules of the game can be used in adaptive dynamics to predict the evolution of the system [15, 16, 19]. Initially, we assume a monomorphic population, with all individuals having the same production and neighborhood size. As is discussed further in Appendix C, in the deterministic limit, the model simplifies to a one-dimensional selection gradient in this case, due to the fact that, if every individual in the population has the same production, each individual gives exactly what it gets in return from its neighborhood, regardless of neighborhood size. Then, the movement of the monomorphic population is governed by the selection gradient in the production direction. As Dieckmann shows, the selection gradient is given by the derivative of fitness with respect to a hypothetical mutant’s trait, evaluated at the trait of the population. With mutant traits (*y*, *n_y_*) and monomorphic traits (*x*, *n_x_*), the production gradient is:

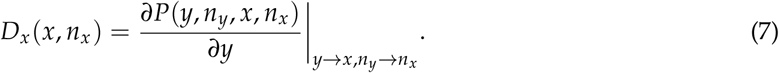

For our model specifically, this production gradient is given by

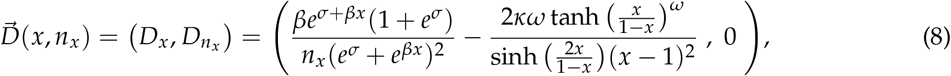

where, as was discussed, the neighborhood size-gradient is zero. Then, we can find the equilibrium points by finding where the selection gradient is zero. For our model, this is a curve in trait space, and can be written so that neighborhood size is a function of production, as follows:

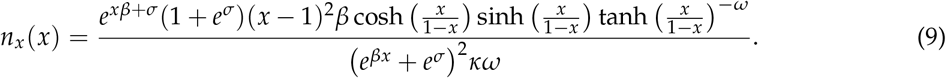

These points can either be attractors of the monomorphic population, or repellors. This can be found in a fairly straightforward way by using the second derivative test to determine if the points are minima or maxima, as is shown in Appendix C. The curve is shown in Figure 2. The dashed part of the curve repels the population. In contrast, the solid line is an attractor. Once the population approaches the stable equilibrium line, it moves along the curve and rich dynamics are possible. This is the focus of the next sections.

### 3.2 Populations along the branching curve favor deterministic branching via saddle instabilit

We want to know what happens when the monomorphic population reaches the attracting set. In this case, it is helpful to think of the individual’s effect on the population’s overall fitness as negligible. While this assumption breaks down with smaller populations, it is helpful for the development of a deterministic approach. So, for the rest of this subsection, we will assume that the individual is an infinitesimal fraction of the population, and so does not affect the mean traits of the population.

Now, the individual(s) may benefit from moving away from the monomorphic population, even if the population won’t benefit from following the individual(s). This process has been called “disruptive selection”, and forces evolution away from the center, toward the extremes in trait space [48]. Finding whether we have disruptive selection is a subtly different calculation than the monomorphic analysis. What we know from the selection gradient calculation is that the population as a whole will not move in any direction. If it did, it would be forced back to the original trait because the curve is attracting.

In order to calculate whether the individual fitness will start increasing as we make a finite move away from the monomorphic population, we take the second derivative, the first nonzero term. Instead of taking the derivative of the payoff function and then setting *y* = *x* and *n_y_* = *n_x_* before taking the next derivative (which would give us the stability of the population), here we care about the individual. This means we take the second derivative and only then do we set *y* = *x* and *n_y_* = *n_x_*.

Unlike the one-dimensional case where points can either be stable or unstable, two- and higher-dimensional trait spaces can have saddle points. If we have a minimum or a saddle, the point is unstable and branching is possible. If the equilibrium is a maximum of individual fitness, the population remains at the equilibrium point, which is a stable strategy.

The second partial derivative test is given by

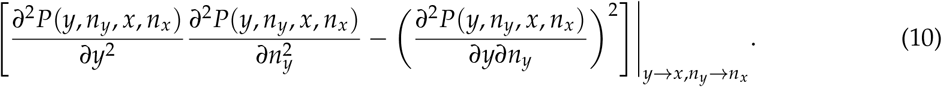

In our case, the second derivative with respect to neighborhood size is zero. Therefore, as long as the cross terms are nonzero, which they are, we have a saddle at every point along the attractor. Therefore, the analytical approach of adaptive dynamics predicts branching as soon as the monomorphic population reaches any part of the attractor set, which we call the branching curve.

### 3.3 Small population size affects the probability of branching and its aftermath

Prototypical results of the game are shown in Figure 2. Multiple possible outcomes can occur for identical simulations with 200 individuals. We can classify all simulations into three scenarios: sustained branching (Figure 2 A), branching followed by extinction of a subpopulation (Figure 2 C), and no branching (Figure 2 E). Figures 2 B, D, F show the corresponding levels of production over time. These results highlight the probabilistic nature of the branching process. By definition, this process cannot be adequately captured by the deterministic approach.

In section 3.2, we showed that in the deterministic limit, our population game always leads to branching (see also Appendix C). In smaller populations, stochastic dynamics play a large role. For our choice of parameter values, motivated by the fact that under these conditions we expect to see coexistence [37] (Table 1), we show that three scenarios are possible. First, we observe branching and coexistence of defectors and cooperators (Figure 2 A,B), Second, we see branching and defector extinction (Figure 2 C,D). Third, we see a scenario of no branching, which then leads to cooperators only (Figure 2 E,F). Note that in this third case, defectors never arise, thus can not go extinct. In the following, we discuss a probabilistic approach to better understand these scenarios. This approach allows us to further understand the branching and extinction processes under stochastic fluctuations in small populations.

### 3.4 Numerical branching and extinction results

Despite the deterministic prediction that branching should occur, many of the simulated populations fail to branch. There can be qualitatively different behaviors which are inherently probabilistic, as seen in Figure 2. We ran multiple simulations to quantify this behavior. The results give us the probability of branching at any given population size, shown in Figure 3 A. Additionally, in Figure 3 B, we find the rate of extinction for a given population size, by counting the generations from branching to extinction, if it occurs. Specifically, we begin counting for extinction when the cooperator subpopulation reaches the full cooperation state of maximum production and minimum neighborhood size.

**Figure 3:**
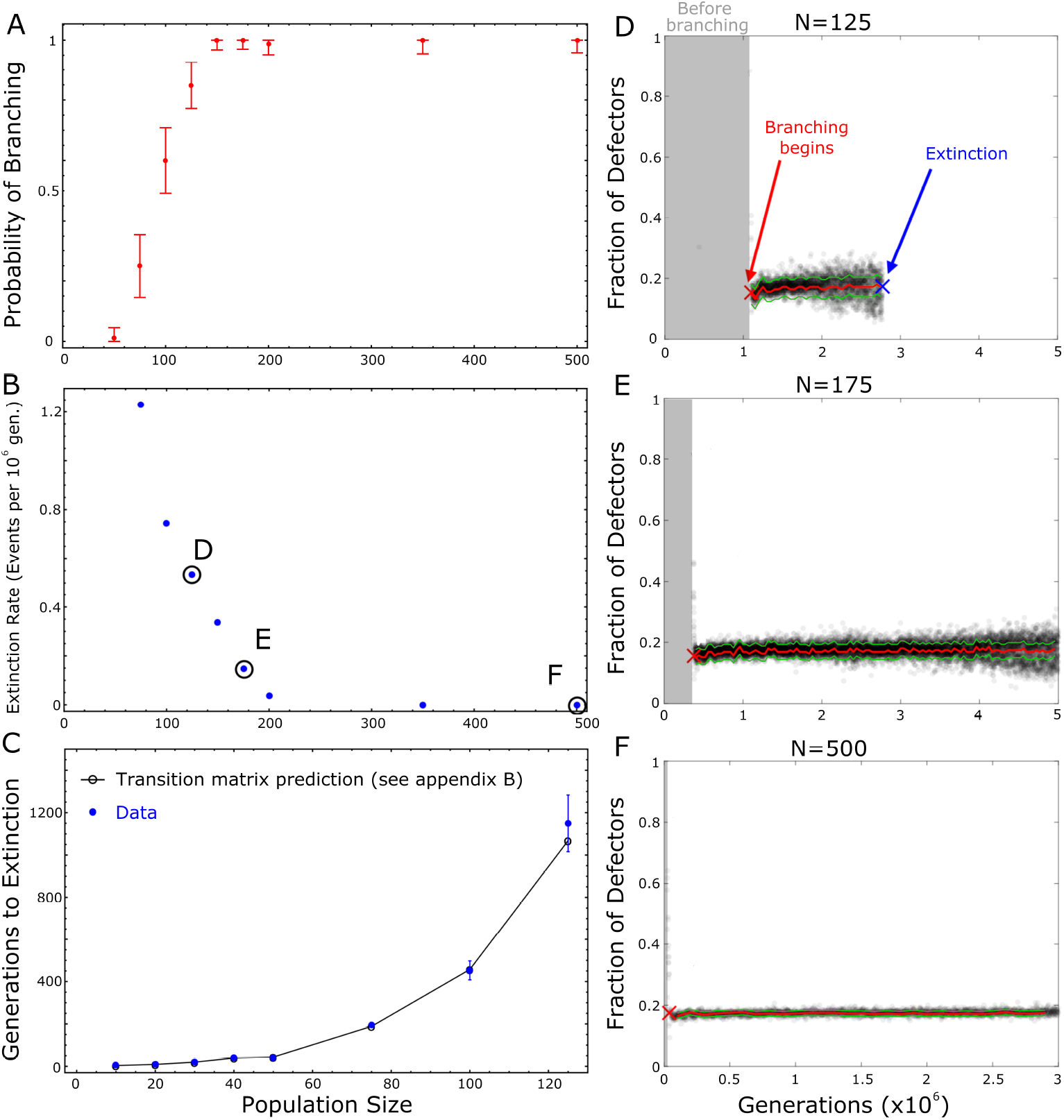
Extinction and branching events strongly depend on population size, N. 100 simulations used for **A-C** except for *N* = 350 and *N* = 500, where only 60 were used. **A:** Branching Probabilities for various population sizes. The calculation of 95% confidence interval via the Agresti-Coull interval method [49] **B:** Extinction Rate for various population sizes, given in the expected number of extinction events occurring in one million generations. Extinction in this case requires branching to occur first. Black circles denote example populations shown in **D-F**. **C:** Extinction time if traits cannot mutate. In this case, we set two initial populations at minimum and maximum allowed traits (full cooperators and defectors), with no mutations. Extinction occurred even if branching would not have happened first. The transition matrix approach is discussed in Section 3.5 and detailed further in Appendix D. Error bars for data represent the standard deviation of the mean. 300 simulations used for *N* = 10 80. 100 simulations for *N* = 100 and *N* = 125. **D-F:** Frequency of defectors for one simulation at *N* = 125, 175, 500, respectively. Gray shaded area indicates generations where the population was still monomorphic (e.g. before branching has occurred). Branching begins at red ‘X’. Red line indicates equilibrium frequency (*p*\*) based on traits (see Appendix E). Green indicates plus and minus one standard deviation based on simulated data. Extinction occurs at blue ‘X’ only for **D**.

When we disregard a main assumption of the adaptive dynamics approach, that of sufficiently large population size, we would not expect it to capture the whole process. Indeed, we know that a lower limit at which branching is no longer favorable exists. The trivial choice when there is only one individual cannot branch. However, these results indicate that the inability to branch occurs for populations exceeding 100 individuals.

From this analysis, we can draw some conclusions, and also suggests some possible hypotheses. We can conclude that, in small populations, a simple adaptive dynamics approach may fail. However, in line with Wakano and Iwasa, our results seem to indicate three distinct possibilities; one where branching is impossible, one where branching is stochastic, and one where branching is deterministic [39]. In the case of stochastic branching, the population may, after reaching a singular point, remain monomorphic for a period of time before branching, or, as mentioned in section 2.2, may miss the opportunity to branch. Deterministic branching, on the other hand, occurs almost immediately after the population reaches a singular point.

### 3.5 Post-branching adaptive dynamics and the cause of defector extinction

Our numerical results suggest the possibility that extinction and branching are related. Therefore, we sought to determine why extinction occurs and under which scenarios it is favored. We were able to identify two possibilities; either extinction occurs because mutational drift moves a subpopulation into a region of trait space unfavorable for coexistence (fitness-based extinction), or stochastic fluctuations in subpopulation frequency away from the mean lead to an absorbing state despite the fact that the subpopulation is in a favorable region for coexistence (frequency-based extinction). While adaptive dynamics may not perfectly capture the extinction and branching possibilities, it still makes reasonable predictions on the mean frequencies of the subpopulations. We use the data from our *k*-means analysis, and expand upon the discussion of subpopulation frequency in Doebeli et al., to find an expected frequency which will allow us to determine whether a subpopulation was located in a “favorable region for coexistence” [19]. In order to do this, we find the frequency of free-riders at which the payoff of a free-rider was equal to that of a producer. The frequency of free-riders that satisfies this will be denoted *p*\*. The details of this calculation are shown in full in Appendix E.

Given *p*\*, we can determine the primary cause of extinction. The expected frequency of free-riders (see Appendix E) is shown as the red line in Figures 3 D-F. In the smaller population size, we see that the variance about that mean is greatly increased, but it nonetheless is centered on the predicted frequency. One of the two possibilities for extinction is that the traits mutate and become unfavorable, leading to extinction based solely on fitness, which we call fitness-based extinction. This would be indicated by the red line going to zero in Figure 3. Clearly, this does not occur because the free-rider frequency is stable (red line), showing only minor fluctuations at the beginning and eventually leveling off to a constant value near 0.17. Therefore, extinction must occur due to stochastic fluctuations in subpopulation frequency causing it to hit the absorbing state, which in this case is a state of all producers. This extinction can have significant impacts on the overall evolution of the system, as is the case for our model. As shown in Figure 2, extinction leads to an all cooperator state, which is an evolutionary stable strategy and is not viable to branch once again. This finding also allows us to predict the time to extinction in certain cases using a transition matrix, which is discussed below.

To investigate the time to extinction, we employ a few simplifying assumptions and utilize a transition matrix approach, shown in Figure 3 C. These assumptions required removing mutations and seeding an already dimorphic population, one subpopulation at maximum production and minimum neighborhood size (cooperators) and one subpopulation at minimum production and maximum neighborhood size (defectors). The mathematical details are shown in Appendix D. All the individuals are updated at each new generation. Therefore, if there is a certain number of defectors in the current generation, it is possible that there would be zero in the next generation. This would only require the chance that all the defectors lose to their opponents. Immediately it becomes clear that extinction is not impossible for larger populations, but becomes increasing unlikely due to the number of defectors that would need to lose in order for extinction to occur.

We can also extend the post-branching adaptive dynamics from Doebeli to two dimensions, creating a vector field which tells us where the defector subpopulation will move [19]. This result is shown in Figure 4. An overlay of the resulting vector field, the details of which is outlined in Appendix E, shows the trait movement of simulations approximates the predicted movement according to our adaptive dynamics approach. These results show a population of 150 individuals, with a defector subpopulation of less than 30 individuals, and is therefore subject to the stochastic effects of small populations. The vector-field outcome, as well as the lack of drift-induced extinction, indicates that adaptive dynamics is still successful in predicting trait space evolution in small populations, even if probabilistic methods are more successful in addressing stochastic fluctuations in subpopulation frequency.

**Figure 4:**
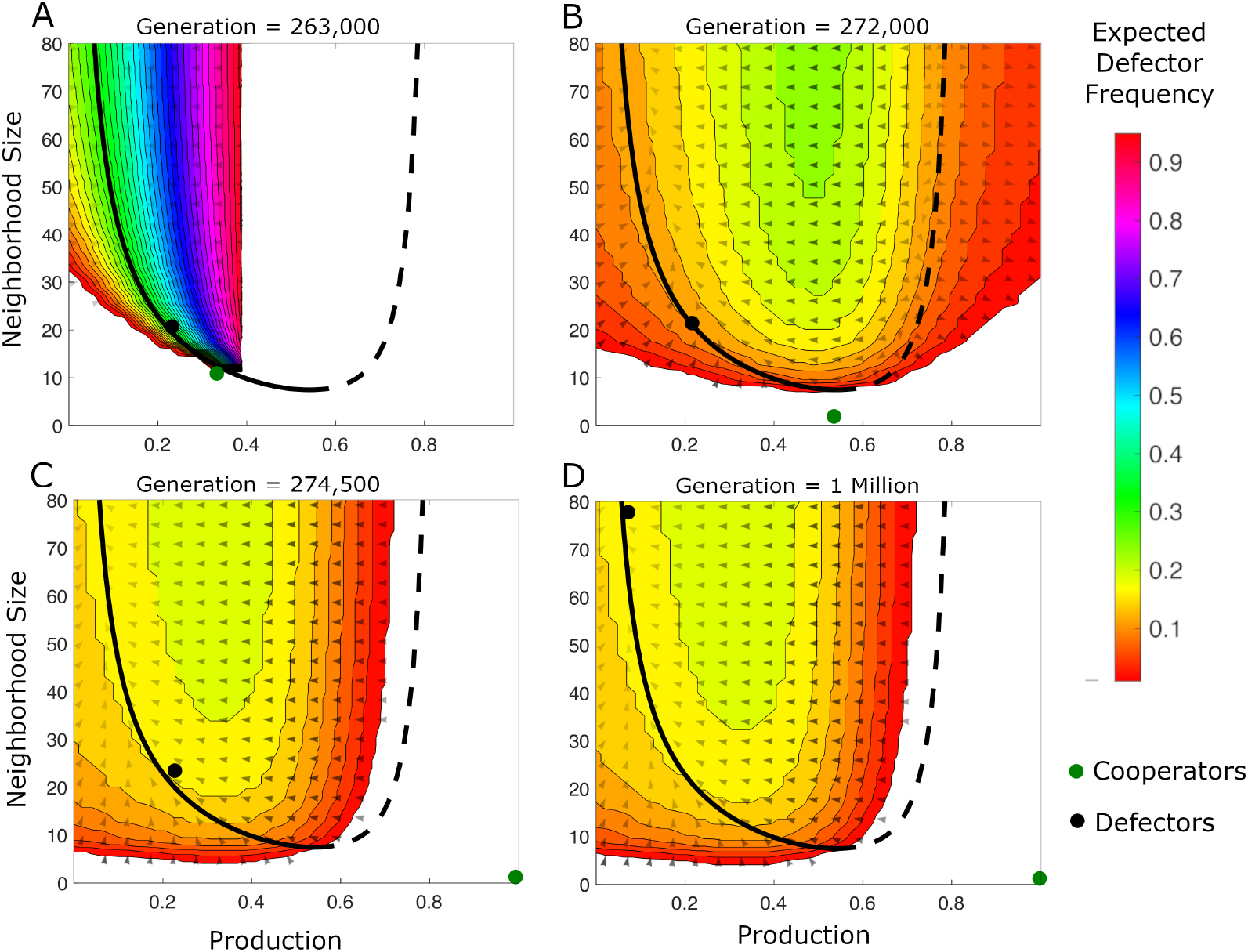
Adaptive Dynamics captures trait evolution. The green dot represents the cooperator subpopulation at the given generation in a simulation of 150 individuals, while black dot indicates defector subpopulation. Filled contour region indicates where the defectors can exist in the deterministic limit, with the color indicating their equilibrium frequency. Blank region indicates where heterogeneity cannot exist, because either defectors (*p*\* > 1) or cooperators (*p*\* < 0) are expected to dominate based on the equilibrium frequency (see Appendix E). Arrows indicate the vector field, with opacity denoting magnitude. The solid and dashed black line indicates the equilibrium points at which the selection gradient is zero for a monomorphic population. Solid represents the attracting set, and dashed represents the repulsive set, as in Figure 2. **A:** Immediately following branching, we see the subpopulations separate quickly, though the vector field is weak. There is a large span in the equilibrium frequency of the subpopulations. **B:** Defector subpopulation has moved little within the weak field, but cooperators make their way quickly towards full production and minimum neighborhood size. The equilibrium frequency of defectors begins to settle. **C:** Cooperators continue moving towards full production and have roughly reached minimum neighborhood size. **D:** If we let many generations pass, we see that the defector subpopulation responds to the underlying deterministic gradient, and slowly increases its neighborhood size accordingly and settles at low production.

## 4. Discussion

Our results highlight the utility and shortcomings of deterministic adaptive dynamics approaches to explain stochastic populations dynamics. Depending on population size, there is a threshold below which deterministic approaches largely fail to capture interesting events. This threshold is likely trait- and game-dependent and drives the emergence of important phenomena such chance and timing of branching or extinction events. We here examined extent to which this stochasticity impacts evolution, and determined the population size that is large enough to avoid impact of extinction. In future work, such a modeling approach can be used to statistically quantify the effects of the public goods game’s key parameters on branching and extinction.

At present, Wakano and Iwasa [39] have presented the best approach to capture branching in finite populations, and their framework is consistent with our results, matching closely with our findings reported in Figure 4 A. Debarre and Otto [41] came to similar conclusions even after modifying the evolutionary game to allow for a varying population size. Although we kept population size fixed, we extend the adaptive evolutionary game dynamics to two a dimensional trait space, which led to two important findings. First, our simulations support the idea that previous work on finite population branching can be generalized to more than one dimension. Second, we show that there exists a finite window in which branching can occur, leading to possibly much faster simulation approaches due to the introduction of certainty that branching has occurred or will not occur. If the stochastic process does not branch by the time the population moves out of the specified window in which branching is favored, it will never branch.

The finding of a definitive and finite branching window can have important impacts on real populations, which may present with small population sizes and multiple co-dependent traits. If the “window” is missed, the course of evolution in such populations could be changed irreversibly. For this reason, it would be useful to further extend and analyze the work of Wakano and Iwasa to multiple dimensions [39]. As a result, one might be able to predict the probability that certain populations will remain monomorphic, branch, or branch and then go extinct.

Our model also has some potential drawbacks. We model selection as a pairwise comparison between a focal individual and a randomly chosen opponent. A continuous-time birth-death process that attempts to more closely mirror cell replication may shift the results in a significant way. In addition, the extinction we observe is often a result of the discrete generational time step. If defectors automatically became more fit as soon as they were less frequent, extinction might be much less likely. A comparison between these two implementations would be helpful to further our understanding of the impact of small populations.

It would be helpful to know if the effect of small populations could be relevant far beyond the few hundred that we observe. The construction of the neighborhood could play an important role and was not considered in detail here. We assumed that a focal cell randomly chooses its neighbors, but perhaps it can change its size as it searches locally. That is, suppose an individual selects neighbors within a fixed radius. Based on the composition it observes in its neighborhood it may choose to increase or decrease that radius in the next generation depending. This additional adaptive potential would impose a type of directed motion in trait space.

There are also many variations on our game which might produce additional interesting insights. While we implemented a cost in production, there is also the possibility of implementing a cost to neighborhood size. Such a cost could, for example, be thought of as a natural drawback to increased motility or neighborhood sensing. Similarly, it might be interesting to implement an adaptation of our game within the framework of a fixed benefit, but with a shared cost, leading to a multi-player “snowdrift” game.

As we have shown, extinction often occurs due to the stochastic fluctuations about the theoretical equilibrium frequency, *p\**. We also note that this process’s variance drops off as 1/*√N* for larger populations. However, smaller populations seemed to reach an upper limit in their deviation, possibly due to relevant covariances, or due to the selective bias of extinction, because exceeding a certain deviation would cause extinction. We also notice a possible link between extinction and branching. Where branching becomes much more likely, the extinction rate drops quickly to near zero. Perhaps the process that limits branching in the first place is similar to, or the same as, the process which causes extinction. Thus, our work serves to highlight the richness of adaptive dynamics models in the stochastic regime.

## Acknowledgements

The authors would like to thank the Moffitt Summer Undergraduate Program to Advance Research Knowledge (SPARK) for providing a platform that led to early results presented in this manuscript.

## Code availability

The source code for simulations and figure generation is available online: adaptivePGG.

## Appendix

## A Simulation Procedure

First, the initial traits are assigned to the population, drawing from independent, user-specified normal distributions for neighborhood size and production. Then, a neighborhood array is randomly generated at initialization. The neighborhood array will assign each individual a random permutation of other individuals which will be its neighbors, but the individual’s neighborhood size, *n*, determines how many neighbors it interacts with. The individual is counted as its own neighbor, so that the minimum neighborhood size is 1. While *n* is a continuous trait, we round to the nearest integer at each generation to calculate the discrete number of neighbors. Each generation, the starting index in the neighborhood array is randomly selected, and then neighbors are sequentially selected until there are *n* neighbors. This method seeks to replicate random selection of neighbors each generation, without the computational expense. In comparing our method with random permutation of neighbors each generation, the results are indistinguishable.

Once the neighbors have been decided, we can calculate the payoff of each individual. The payoff of the opponent must be greater than the payoff of the individual for the opponent to have a nonzero probability of replacing the individual. If the opponent is more fit than the focal individual, the probability of opponent replacing individual is the payoff difference between focal and opponent normalized by *α*, the maximum payoff difference between any two individuals in that given generation. If the individual is not replaced, the parent replaces itself, and the individual’s traits are moved forward to the next generation, barring any mutation. Mutation is possible regardless of whether the opponent or individual becomes the parent, with probabilities *μ_x_* and *μ_n_*, which were both set to 0.01. Mutations are independent in neighborhood size and production, and are drawn from a normal distribution with parent trait as the mean and user-specified standard deviation. As mentioned previously, the standard deviation of mutation we used was 0.2 for neighborhood size and 0.005 for production. The pseudo-code is listed below.

## Pseudo-code

1. Input parameters and initialize population
2. Initialize randomly sorted array of neighbors for each individual
3. For each generation:

a. Loop through population, assigning payoffs:

i. Assign neighbors based on the individual’s neighborhood size, their array of neighbors, and a random starting index for that array of neighbors.
ii. Calculate payoffs of each individual
b. Loop through population again, determining offspring traits:

i. Select random opponent
ii. Calculate probability of replacement and determine which individual becomes the parent
iii. Assign parent’s traits to next generation
iv. Check for mutations independently in production and neighborhood size

## B K-means Approach to Locate the Branching Point

The input for the *K*-means analysis is the array of production traits for each given generation. Using the MATLAB R2019b ‘kmeans’ function, we force the production array to be separated into two groups. Then, subtracting the mean of one group from the other, we can see how far apart the “groups” are. Beyond a certain threshold, we declare that branching had occurred. We set this threshold to be consistent with what we see in the resulting plots. Specifically, we want to ensure that branching which is called actually appears as a branching event, with two clearly separate groups that maintain separation for several generations. Typically, this threshold is between 0.1 and 0.2, though we use 0.2. With this value, no false positives occurred during our simulations. However, the declaration that a branch had occurred was delayed by a few generations. This shift was very small relative to the total number of generations and consistent among different simulations and so we neglected its impact.

## C Monomorphic Population and Selection Gradient

Given a payoff function with a benefit B and cost C, as shown in Figure 1 C, we want to know where the population is favored to move. For this, we take an adaptive dynamics approach, which begins by predicting the movement in trait space of a monomorphic population. The payoff function we use looks like this:

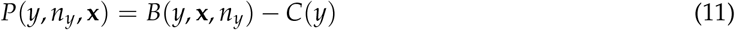

Where *P*(*y*, *n_y_*, **x**) denotes an individual with production *y* and neighborhood size *n_y_*, with neighbors **x**. The neighbors represent a vector, but we assume the population is monomorphic with trait *x*. The form of our payoff function, as shown above, is:

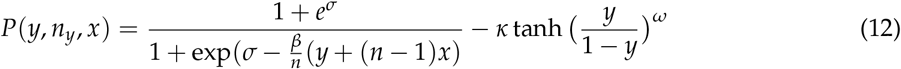

The selection gradient in two dimensions is the gradient of a mutant with trait (*y*, *n_y_*) in a population where all other individuals have trait (*x*, *n_x_*). The population does not necessarily move in the direction of increasing population fitness, but instead moves in the direction of increasing individual fitness. It is intuitive that this should happen because a mutant with higher payoff will eventually replace a resident of lesser fitness. We can re-write our payoff function with a slightly different notation, 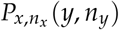, indicating that we are operating on the mutant, *y* and *n_y_*. Then, the selection gradient becomes:

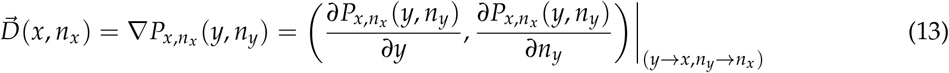

In our case, the vector selection gradient is zero for neighborhood size and reduces to the one-dimensional production derivative. This is because, for our payoff function, if the individual has the same production as the rest of the population, there is no change in the observed production due to a different neighborhood size (each individual is getting back exactly what they put in). Thus, for *y* = *x*, the neighborhood size is irrelevant in determining the movement of the entire population. With the default parameters shown in the model introduction (figure 1B), our selection gradient is:

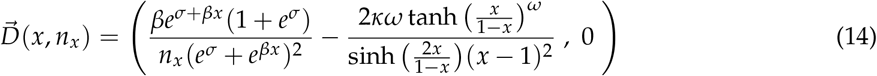

Selection gradient tell us the movement of the monomorphic population. When the selection gradient is zero (i.e. when 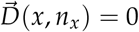 we have singular points. In our case, we find a line of singular points. That is, the selection gradient is equal to zero whenever Eq. 15 is satisfied:

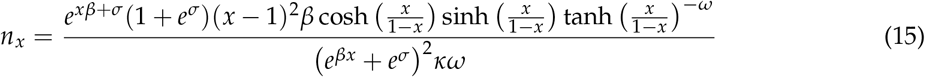

The singular point can either be an attractor (pop. moves towards) or repellor (pop. moves away) of the population as a whole. Physically, we can think of it as a potential well, the gradient is zero but the equilibrium can be either stable or unstable. Generally, this can be found by taking the gradient of the selection gradient, 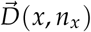, and using the second partial derivative test to determine stability. In our case, however, we have a selection gradient which is only nonzero in one direction. Therefore, while the second partial derivative test would be zero and inconclusive, we can simply use the one-dimensional second derivative test to determine stability:

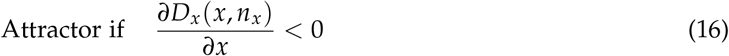

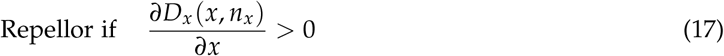

The actual derivative taken for equations 16 and 17 is a few lines long and will not be included. Instead, the figure below shows the curve of singular points and the regions where the points are attractors (solid line) and where they are repellors (dashed line).

The dynamics shown in Figure 5 are fairly simple. The monomorphic population generally moves only in the direction of increasing or decreasing production, depending on where it starts. However, more interesting dynamics can occur along the solid line, once the population stops moving as a monomorphic unit. This is where branching is possible.

**Figure 5:**
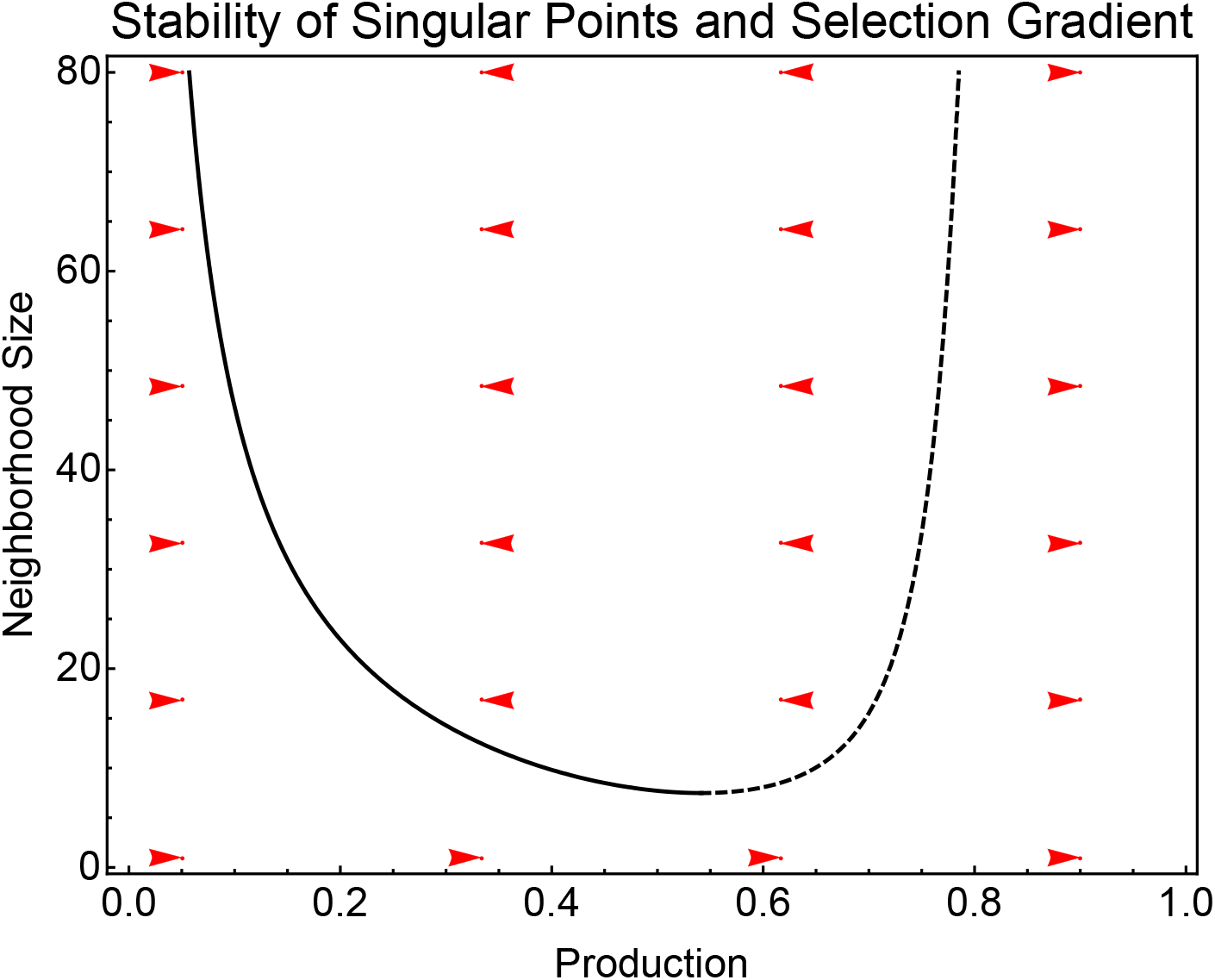
Analysis of the stability of singular points in our model. Attractors are shown as the solid line and repellors are the dashed line. The red vector field represents the selection gradient. As expected, the arrows point towards the solid line and away from the dashed line. Note also that the arrows are all horizontal, indicating a zero gradient in the neighborhood size direction.

## D Transition Matrix Approach to Extinction

Given the competitive process suggested by Doebeli *et al.* [19], which we use in our model, we can employ the following transition matrix to evaluate the probability of extinction of a subpopulation, under these strong assumptions: the population was initialized with two subpopulations, one at full cooperation, with a neighborhood of just itself and maximum production, and the other at zero cooperation, with a neighborhood size equal to population size and no production. We also restricted mutations, the effect of which will be discussed. In this limited form of our model, more fit individuals would always be replaced by less fit individuals, if they chose a more fit opponent. Therefore, the probability of going from one state to another is the probability of choosing the “opponents” that lead to that state.

In a more concrete case, the probability of going from a state with 4 cooperators to a state with 5 cooperators requires two things. First, cooperators must be more fit in order to replace defectors. This is dependent on the frequency of the cooperators. Second, the defectors must select their “opponent” so that only one defector selects a cooperator, loses, and is replaced by a cooperator. Because each defector is equally likely to select any individual (except itself) as an opponent, we can find the probability of a defector choosing a cooperator, which would simply be the number of cooperators divided by *N* − 1, the total number of possible opponents. From there, we can find the binomial distribution telling us the probability of having *X* defectors in state *a_i_*, and having *Y* defectors in state *a*_*i*+1_.

There are two absorbing states in this stochastic matrix, each representing extinction of a subpopulation. In our model, due to the lower frequency of defectors, we only observe extinction of defectors, but this is only dependent on the payoff function we chose, and generally extinction of either subpopulation can occur. If we define *T* as our transition matrix, we can break it down into *Q*, the transient states, and *R*, the absorbing states (a=0 or a=N), as follows:

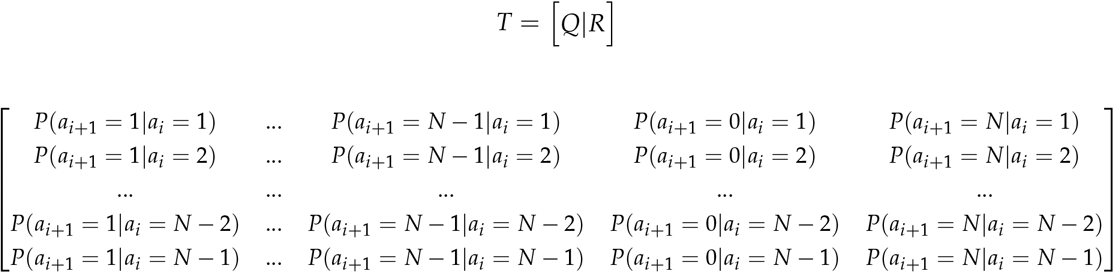

For a population of five individuals, this becomes a bit easier to visualize:

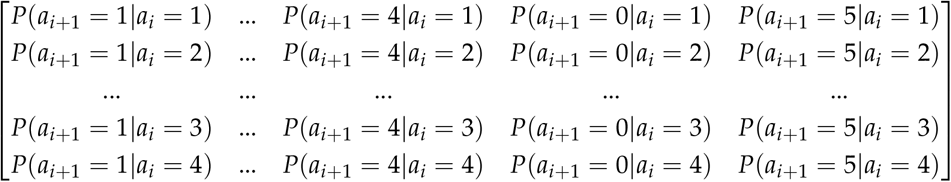

Written out, the probability of going from *m* defectors to *n* defectors in one step is either one of two probabilities. Either defectors are less frequent than equilibrium, in which case they are more fit, or they are more frequent and less fit. In the case where defectors are less frequent, we must gain defectors, or at least stay the same. Valid for *n* ≥ *m* (zero elsewhere), if *m*/*N* < p* (less frequent than equilibrium), we have:

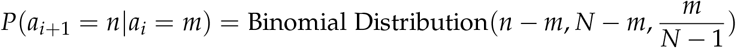

On the other hand, if defectors are over-frequent (*m*/*N* > *p*\*), the probabilities become zero unless *n* ≤ *m*, in which case the probability is:

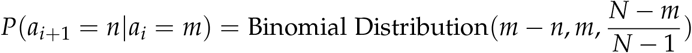

The full numerical matrix, *T*, for five individuals with *p*\* =.1505 then becomes:

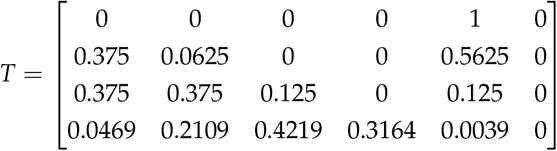

*N*, showing only the transient states, is:

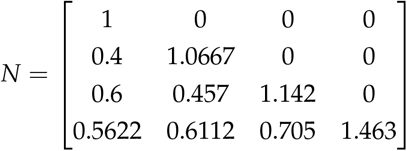

*R*, showing only the absorbing states, is:

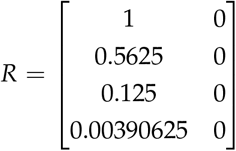

*B*, showing the probabilities of being absorbed by a state given a starting state (row), is:

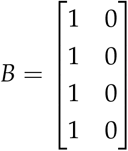

*t*, showing the expected number of steps given a starting state (row), is:

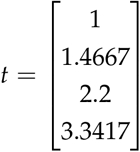

When we extend this to not just 5 individuals, but 75 or 100, we can find the probability of extinction given our restrictions on the model. The results can be seen in Figure 4 C, which shows fairly good agreement in the smaller population sizes. It becomes fairly computationally difficult to run for larger population sizes due to the fact that the number of expected generations until extinction increases so quickly. Still, our success in small *N* allows us to predict this expected value without requiring computationally expensive simulation.

Another interesting issue is the effect of mutations, generally we see that allowing mutations leads to a slight increase in the number of generations the subpopulation survives for. In the case of *N* = 75, the number of generations before extinction jumped from roughly 195 to 253. This is likely due to the model itself, because the replacement probability is determined by the fitness difference. The replacement probability would either be 1 or 0 in the case of no mutation. However, once we allow mutation, and the resulting variation in fitness, a less fit individual facing a more fit individual would have a nonzero survival chance as long as it wasn’t the least fit facing the most fit. This becomes increasingly difficult to incorporate into a transition matrix approximation, so it is likely that our transition matrix results represent a lower value than would be observed with mutation. Nonetheless, it can provide a ballpark estimate in addition to showing that, for increasing *N*, the time to extinction increases quite rapidly (Figure 6).

**Figure 6:**
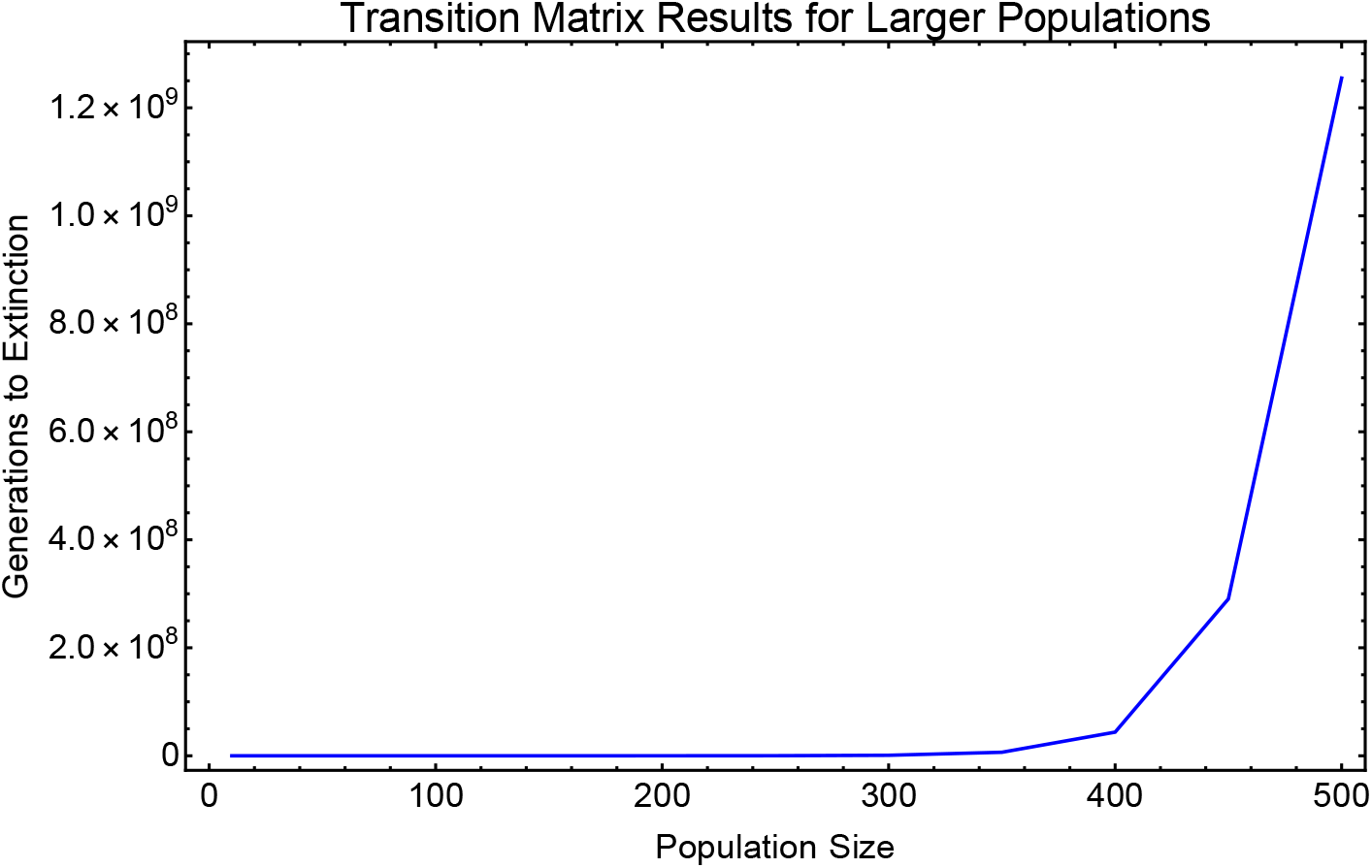
For population sizes up to 500, we see that extinction becomes increasingly unlikely to observe in a reasonable number of generations. Though we cannot simulate for billions of generations, the agreement between transition matrix and simulations at small populations allows us to argue that extinction of a branch is unlikely for large populations.

## E Equilibrium Frequency and Vector Field

In order to find the equilibrium frequency of the two subpopulations we essentially just set the fitnesses equal to each other. In practice, this is fairly simple, but it is slightly more complicated by our neighborhood selection process. In other words, if we simply took the average available public good in our calculation of the fitness, this would be simple. But, we randomly select neighbors, meaning it is possible to have a neighborhood entirely composed of cooperators or entirely composed of defectors. On average, the fraction of neighborhood composed of defectors will be equal to the fraction in the population as a whole, but the nonlinearity of the payoff means we must consider each possible neighborhood. Therefore, expanding on the work in Doebeli et al. [19], the equilibrium frequency (*p*\*) is found by setting the expanded payoff functions equal. For our purposes, we assume a branched population with traits (*x*, *n_x_*) and (*y*, *n_y_*). **P** () denotes the payoff function whereas *P*(*i|n*) denotes probability of selecting *i* neighbors of type *y* given a neighborhood size. It should be noted that for our purposes, all neighborhood sizes (which may take on non-integer values) are rounded to integer values for this calculation and in the simulations:

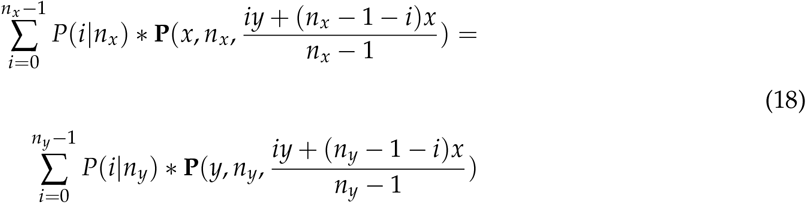

As we see, we can weight the various payoffs resulting from the varied neighborhoods by the probability of generating that neighborhood. This probability is a hypergeometric distribution which depends on the equilibrium frequency, *p*\*, and the population size *N*. In the infinite population size case, this would be a binomial distribution, but with finite population size we must take into account the fact that we are drawing without replacement. Because *i* represents (arbitrarily) the number of *y* individuals chosen for the neighborhood, this probability has slightly different form for the *x* and *y* case. The difference stems from the fact that we cannot chose the focal individual as our neighbor and therefore have one less *y* to choose when *y* is the focal. We let *p*\* represent the equilibrium fraction of defectors. In this case, the probabilities are:

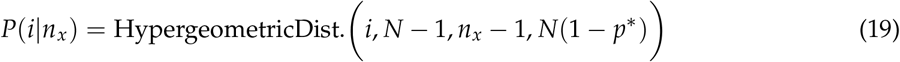

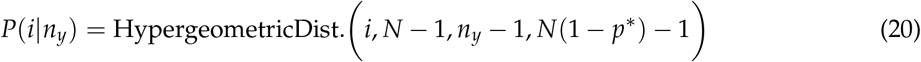

Numerically solving equation 18 for specific *x*, *y*, *n_x_*, and *n_y_* values gives us a value for *p*\*, telling us the expected frequency of the individuals with trait (*x*, *n_x_*). This can be seen as the red lines in Figure 4 D-F as well as in the contours in Figure 3.

Using this calculation of *p*\* also allows us to also find the direction of expected movement of the population. Similar to the one-dimensional case, we want to know what what direction a mutant must mutate to in order to be more fit than the “resident”. This becomes slightly more complicated by the fact that there are two subpopulations, but otherwise it is fairly straightforward. Again, this originates from the work of Doebeli, and we extend it to two dimensions. First, we must find the invasion fitness of a mutant with traits (*v*, *n_v_*). This involves calculating its fitness in addition to subtracting from the average fitness of the population, 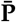. 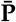 can be given simply by the payoff of a cooperator or defector at the frequency *p*\*, because we know these are set to be equal. Then, we must find the payoff of the traits (*v*, *n_v_*) in order to find the invasion fitness **P** (*v*, *n_v_*):

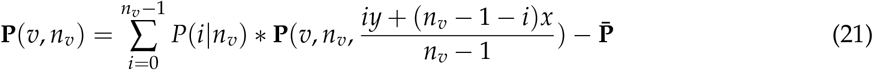

If we take the gradient of this invasion fitness, evaluated at one of the subpopulations ((*x*,*n_x_*) or (*y*, *n_y_*)), we have the vector field indicating the movement of each subpopulation in trait space. This is shown for the defector subpopulation in Figure 3.

There are two small but important details pertaining to this vector field. First, because neighborhood size is converted to integer values, we don’t take an infinitesimal derivative with respect to neighborhood size. Instead, we increment by 1 to evaluate a more grainy, but still useful, derivative. Second, to account for the wide variation in trait magnitude, we had to normalize by the step size of mutations. Because mutation in neighborhood size was 40 times greater than mutation in production, the gradient was adjusted by multiplying the neighborhood size derivative by 40. The result, shown in Figure 4, largely matches the simulations, though random drift is also relevant, and more so at smaller population sizes.

## References

1. Rood JP. Population dynamics and food habits of the banded mongoose. African Journal of Ecology. 1975;13(2):89–111.

2. Zomorrodi AR, Segrè D. Genome-driven evolutionary game theory helps understand the rise of metabolic interdependencies in microbial communities. Nature Communications. 2017;8(1):1563.

3. Frank SA. Foundations of social evolution. Princeton: Princeton Univ. Press; 1998.

4. Archetti M, Ferraro DA, Christofori G. Heterogeneity for IGF-II production maintained by public goods dynamics in neuroendocrine pancreatic cancer. Proceedings of the National Academy of Sciences USA. 2015;112:1833–1838.

5. Sober E, Wilson DS. Unto Others: The Evolution and Psychology of Unselfish Behavior. Cambridge, MA: Harvard University Press; 1998.

6. Wilson DS, O’Brien DT, Sesma A. Human prosociality from an evolutionary perspective: variation and correlations at a city-wide scale. Evolution and Human Behavior. 2009;30.

7. Fitzpatrick BM, Fordyce JA, Gavrilets S. What, if anything, is sympatric speciation? Journal of Evolutionary Biology. 2008;21.

8. Bolnick DI, Fitzpatrick BM. Sympatric Speciation: Models and Empirical Evidence. Annual Review of Ecology, Evolution, and Systematics. 2007;38.

9. Swanton C, Burrell RA, Futreal PA. Breast cancer genome heterogeneity: a challenge to personalised medicine? Breast Cancer Research. 2011;13.

10. Turner NC, Reis-Filho JS. Genetic heterogeneity and cancer drug resistance. The Lancet Oncology. 2012;13.

11. McGranahan N, Swanton C. Biological and Therapeutic Impact of Intratumor Heterogeneity in Cancer Evolution. Cancer Cell. 2015;27.

12. Turajlic S, Sottoriva A, Graham T, Swanton C. Resolving genetic heterogeneity in cancer. Nature Reviews Genetics. 2019;20:404–416.

13. Hofbauer J, Sigmund K. Evolutionary Games and Population Dynamics. Cambridge University Press, Cambridge; 1998.

14. Geritz SA, Metz JA, Kisdi É, Meszéna G. Dynamics of adaptation and evolutionary branching. Physical Review Letters. 1997;78(10):2024.

15. Diekmann O. A beginner’s guide to adaptive dynamics. Mathematical Modelling of Population Dynamics. 2003;63.

16. Champagnat N, Ferriere R, Arous GB. The canonical equation of adaptive dynamics: a mathematical view. Selection. 2001;2.

17. Guttal V, Couzin ID. Social interactions, information use, and the evolution of collective migration. PNAS. 2010;107.

18. Doebeli M. A model for the evolutionary dynamics of cross-feeding polymorphisms in microorganisms. Population Ecology. 2002;44.

19. Doebeli M, Hauert C, Killingback T. The evolutionary origin of cooperators and defectors. Science. 2004 Oct;306(5697):859–62.

20. Hardin G. The tragedy of the commons. Science. 1968;162:1243–1248.

21. Hardin G. Extensions of “The Tragedy of the Commons”. Science. 1998;280:682–683.

22. Pepper JW. Drugs that target pathogen public goods are robust against evolved drug resistance. Evolutionary Applications. 2012;5(7):757–761.

23. Davidson AD, Lightfoot DC. Keystone rodent interactions: prairie dogs and kangaroo rats structure the biotic composition of a desertified grassland. Ecography. 2006;29(5):755–765.

24. Dugatkin LA. Tendency to inspect predators predicts mortality risk in the guppy (Poecilia reticulata). Behavioral Ecology. 1992;3(2):124–127.

25. Whitehouse ME, Lubin Y. The functions of societies and the evolution of group living: spider societies as a test case. Biological Reviews. 2005;80(3):347–361.

26. Drescher K, Nadell CD, Stone HA, Wingreen NS, Bassler BL. Solutions to the public goods dilemma in bacterial biofilms. Current Biology. 2014;24(1):50–55.

27. Nadell CD, Bassler BL, Levin SA. Observing bacteria through the lens of social evolution. Journal of Biology. 2008;7(7):27.

28. Gore J, Youk H, van Oudenaarden A. Snowdrift game dynamics and facultative cheating in yeast. Nature. 2009;459:253–256.

29. Metz JAJ, Geritz SAH, Meszena G, Jacobs FJA, van Heerwaarden JS. Adaptive dynamics: a geometrical study of the consequences of nearly faithful replication. In: van Strien SJ, Verduyn Lunel SM, editors. Stochastic and Spatial Structures of Dynamical Systems. Amsterdam: North Holland; 1996. p. 183–231.

30. Archetti M, Scheuring I. Evolution of optimal Hill coefficients in nonlinear public goods games. Journal of Theoretical Biology. 2016;406(7).

31. Archetti M, Scheuring I. Review: Game theory of public goods in one-shot social dilemmas without assortment. Journal of Theoretical Biology. 2012;299(21):9–20.

32. Hauert C, Michor F, Nowak MA, Doebeli M. Synergy and discounting of cooperation in social dilemmas. Journal of Theoretical Biology. 2006;239:195–202.

33. Cornforth DM, Sumpter DJ, Brown SP, Brännström Å. Synergy and group size in microbial cooperation. The American Naturalist. 2012;180(3):296–305.

34. Gerlee P, Altrock PM. Complexity and stability in growing cancer cell populations. Proceedings of the National Academy of Sciences USA. 2015;112:E2742–E2743.

35. Gerlee P, Altrock PM. Extinction rates in tumour public goods games. Journal of The Royal Society Interface. 2017;14(134):20170342.

36. Gerlee P, Altrock PM. Persistence of cooperation in diffusive public goods games. Physical Review E. 2019;99(6):062412.

37. Kimmel GJ, Gerlee P, Brown JS, Altrock PM. Neighborhood size-effects shape growing population dynamics in evolutionary public goods games. Communications Biology. 2019;2(53).

38. Kimmel GJ, Gerlee P, Altrock PM. Time scales and wave formation in non-linear spatial public goods games. PLoS computational biology. 2019;15(9):e1007361.

39. Wakano JY, Iwasa Y. Evolutionary Branching in a Finite Population: Deterministic Branching vs. Stochastic Branching. Genetics. 2013;193.

40. Claessen D, Andersson J, Persson L, de Roos AM. Delayed evolutionary branching in small populations. Evolutionary Ecology Research. 2007;9.

41. Debarre F, Otto SP. Evolutionary Dynamics of a Quantitative Trait in a Finite Asexual Population. Theoretical Population Biology. 2016;108.

42. Isaac RM, Walker JM, Williams AW. Group size and the voluntary provision of public goods: Experimental evidence utilizing large groups. Journal of Public Economics. 1994;54(1):1–36.

43. Driscoll WW, Pepper JW. Theory for the evolution of diffusible external goods. Evolution. 2010;64(9):2682–2687.

44. Archetti M, Scheuring I. Review: Evolution of cooperation in one-shot social dilemmas without assortment. Journal of Theoretical Biology. 2012;299:9–20.

45. Constable GW, Rogers T, McKane AJ, Tarnita CE. Demographic noise can reverse the direction of deterministic selection. Proceedings of the National Academy of Sciences. 2016;113(32):E4745–E4754.

46. Brown JS, Vincent TL. Evolution of cooperation with shared costs and benefits. Proceedings of the Royal Society of London B: Biological Sciences. 2008;275(1646):1985–1994.

47. Archetti M. Dynamics of growth factor production in monolayers of cancer cells and evolution of resistance to anticancer therapies. Evolutionary applications. 2013;6(8):1146–1159.

48. Mather K. Polymorphism as an Outcome of Disruptive Selection. Evolution. 1955;9.

49. Agresti A, Coull BA. Approximate Is Better than ‘Exact’ for Interval Estimation of Binomial Proportions. The American Statistician. 1998;52(2).

